# Circulating Myeloid-Derived Suppressor Cell load and disease severity are associated to an enhanced oligodendroglial production in a murine model of multiple sclerosis

**DOI:** 10.1101/2024.07.18.604171

**Authors:** Mari Paz Serrano-Regal, Celia Camacho-Toledano, Inmaculada Alonso-García, María Cristina Ortega, Isabel Machín-Díaz, Rafael Lebrón-Galán, Jennifer García-Arocha, Leticia Calahorra, Diego Clemente

## Abstract

**Background:** Multiple sclerosis (MS) is a chronic, inflammatory and demyelinating disease of the central nervous system (CNS) that is highly heterogeneous in terms of disease severity and tissue damage extent. Improving myelin restoration is essential to prevent neurodegeneration and the associated disability in MS patients. However, remyelinating therapies are failing in clinical trials, in part, due to the absence of classifying biomarkers of different endogenous regenerative capacities amongst enrolled patients. We previously reported that circulating monocytic myeloid-derived suppressor cells (M-MDSCs) at the onset of the murine model of MS experimental autoimmune encephalomyelitis (EAE) are associated with milder disease courses and less degree of demyelination and axonal damage in spinal cord lesions, while at peak are indicative of a better symptom recovery. Moreover, M-MDSCs are able to promote *in vitro* oligodendrocyte precursor cell (OPC) proliferation and differentiation towards mature oligodendrocytes (OLs) through the release of the soluble factor osteopontin.

**Results:** Here, we show a relationship between disease severity and a gradient of OPCs between the rim and the core in mixed active-inactive lesions of MS patients, along with a positive correlation between M-MDSC density and OPC abundance in the same lesions. We also show that EAE disease severity negatively influences the density of total and newly generated OPCs found associated to the demyelinated lesions of the spinal cord at the peak of the disease. In addition, disease severity also impacts the abundance of newly generated OLs originated either during the effector phase or during the early recovery phase. We also demonstrate the positive association between infiltrated M-MDSCs and the abundance of OPCs in the periplaque of demyelinating lesions at the peak of EAE. Interestingly, circulating M-MDSCs at EAE onset and peak of the disease are directly associated to a higher density of newly generated OLs in the plaque and periplaque, respectively.

**Conclusion:** Disease severity clearly impacts oligodendrocyte generation during a neuroinflammatory insult like EAE. Our results set the basis for further studies on M-MDSCs as a promising new biomarker that identify a CNS prone to the generation of new OLs that may contribute to restore myelin.

## BACKGROUND

Multiple sclerosis (MS) is an immune-mediated disease of the central nervous system (CNS) characterized by focal demyelination, gliosis and axonal degeneration that represents the leading cause of non-traumatic disability among young adults [1,2]. The most common form of MS is relapsing-remitting MS (RRMS), which is characterized by episodes of neurological dysfunction (relapses) followed by periods of full or partial recovery (remission; [3]). RRMS is highly heterogeneous in terms of disease course and symptoms [4] and the evolution of MS seems to be the result of the interplay between acute inflammatory relapses, chronic inflammation in the CNS, the degeneration of axons and loss of oligodendrocytes. Increased resilience in the rates of both axon degeneration and the efficiency of remyelination at early stages of the disease have been proposed to be the basis of the heterogeneous severity found in this clinical MS subtype [5].

In MS, axonal loss - and the subsequent neurodegeneration - is directly associated with the permanent functional deficits observed in patients, and it probably originates as a consequence of the vulnerability of denuded axons [6]. The loss of myelin can be repaired by the spontaneous generation of new myelin sheaths around the axons, i.e. remyelination. This process is associated with axonal preservation [6] and lower levels of disability in patients with MS [7]. However, its efficiency decreases significantly with age and disease progression [8]. In this regard, it has been hypothesized that impaired oligodendrocyte precursor cell (OPC) activation, recruitment and differentiation are among the main putative causes of remyelination failure [9–12]. Therefore, improving OPC proliferation/recruitment to demyelinated areas as well as OPC differentiation, could suffice myelinating cells to accomplish effective remyelination and, lately, prevent neurodegeneration. Importantly, interindividual variability in the endogenous ability to induce both processes (proliferation and differentiation of OPCs) in the CNS of MS patients has not been explored to date.

Currently, all of the available disease-modifying treatments for MS are immunomodulators and/or immunosuppressants that reduce disease activity and the frequency of relapses but do not efficiently prevent the progression of disability. Hence, one of the therapeutic challenges in MS is the search for molecules that enhance remyelination and promote neuroprotection, which in turn would stop or slow disease progression [13]. The lack of suitable biomarkers of regeneration together with the fact that there are heterogeneous profiles of remyelination among MS patients [13] could explain, at least in part, why myelin repair molecules are failing in clinical trials. Therefore, the identification of new classifying biomarkers indicative of a CNS prone to efficiently induce OPC proliferation and differentiation would improve the success rate of clinical trials with pro-remyelinating molecules.

Experimental autoimmune encephalomyelitis (EAE), the most commonly used animal model of MS, has been widely employed to study the pathogenesis of the disease as well as to carry out preclinical evaluation of candidate pro-remyelinating molecules [13–17]. Individualized evaluation of EAE appears as an excellent strategy to find clinically translatable biomarkers of responsiveness and efficacy of DMTs [18]. In this sense, this strategy has been used to study the interplay between disease severity and CNS-related histopathological events, as milder disease courses are associated with less demyelination and axonal damage [19]. However, the impact of the heterogeneous disease severity on the ability of the CNS to produce new mature oligodendrocytes (OLs) after a neuroinflammatory insult remains elusive.

Monocytic myeloid-derived suppressor cells (M-MDSCs) are a heterogeneous population of regulatory cells of the innate immune response originated in the bone marrow. Under pathological conditions or inflammatory contexts, they remain undifferentiated and acquire immunosuppressive functions [20]. The presence and characterization of M-MDSCs in the spinal cord of EAE mice at the peak of the disease course was previously described [21]. Interestingly, there is a direct association between the presence of M-MDSCs in the plaque and the presence of OPCs in the adjacent periplaque in spinal cord demyelinating lesions of mice with EAE. Moreover, M-MDSCs influence different aspects of OPC biology *in vitro* (survival, proliferation and differentiation) through the release of the soluble factor osteopontin [22]. Our group has pioneered the morphological demonstration of cells exhibiting all the bona-fide markers of M-MDSCs in *post-mortem* tissue from MS patients. Interestingly, there is a direct relation between the density of M-MDSCs in MS tissue and disease severity [23]. Regarding the EAE model, circulating M-MDSCs at disease onset were associated with milder disease courses and less degree of demyelination and axonal damage in the spinal cord, while at peak were indicative of a better symptom recovery [23]. These results indicate that M-MDSCs are an excellent predictive biomarker of disease severity in the EAE model. However, little is known about whether the abundance of circulating M-MDSCs could predict a better CNS ability to produce new OPCs able to differentiate into OLs.

In this work, we found a direct association between disease duration and the relative density of OPCs in mixed active/inactive lesions (mAIL) of MS patients. Moreover, the density of OPCs at the rim of these lesions (rAIL) directly correlated with the abundance of M-MDSCs analyzed in the same areas. On the other hand, we demonstrate that milder courses of EAE are associated with a higher presence of proliferating OPCs at the end of the effector phase and of newly generated OLs at the end of the recovery phase, in both cases in the periplaque of demyelinated spinal cord lesions. Notably, a higher content of circulating M-MDSCs at the onset or the peak of the disease is associated with a plaque and periplaque enriched in newly generated OLs, respectively, once each animal reached its maximal clinical recovery. These results pave the way for the potential use of circulating M-MDSCs as a promising biomarker of a CNS more prone to produce new OLs in inflammatory conditions such as MS.

## RESULTS

### Disease duration is associated to the presence of an OPC gradient in mAIL of MS patients

Previous reports have shown that the density of OPCs varies between different regions of demyelinating plaques and the surrounding periplaque and normal appearing white matter (NAWM), finding higher or lower numbers than in the control white matter [24]. In order to investigate whether disease severity impacts on the relative abundance of OPCs associated to the different inflammatory environments of MS lesions, we analyzed the core (cAIL), rim (rAIL) and NAWM of mAILs from 8 MS patients with different disease durations of their clinical course (Table 1). We found that the abundance of OPCs was lower in the cAIL than in the rAIL and adjacent NAWM, with no differences detected between the latter two (Figure 1A-E). As expected, disease duration was independent of the ratio between the adjacent NAWM and the rAIL (r = - 0.405, p = 0.290). Interestingly, we found a positive association between the disease duration and the ratio of the densities of OPCs between the rAIL and the cAIL of each analyzed patient (Figure 1b’-b’’, d’-d’’, F). We previously reported a direct relationship between the density of M-MDSCs and OPCs in the spinal cord demyelinated lesions of mice with EAE [22]. Within mAIL, M-MDSCs were mainly present in the rAIL in progressive MS patients [23]. Of note, the higher the density of M-MDSCs, the higher the abundance of OPCs in the rAIL (Figure 1G). Overall, these results indicate that disease severity negatively affects the relative abundance of OPCs present in the different areas of mAIL from MS patients. Conversely, the greater presence of the immunoregulatory M-MDSCs in the rAIL was positively related to an enrichment of OPCs in the same area.

**Figure 1.**
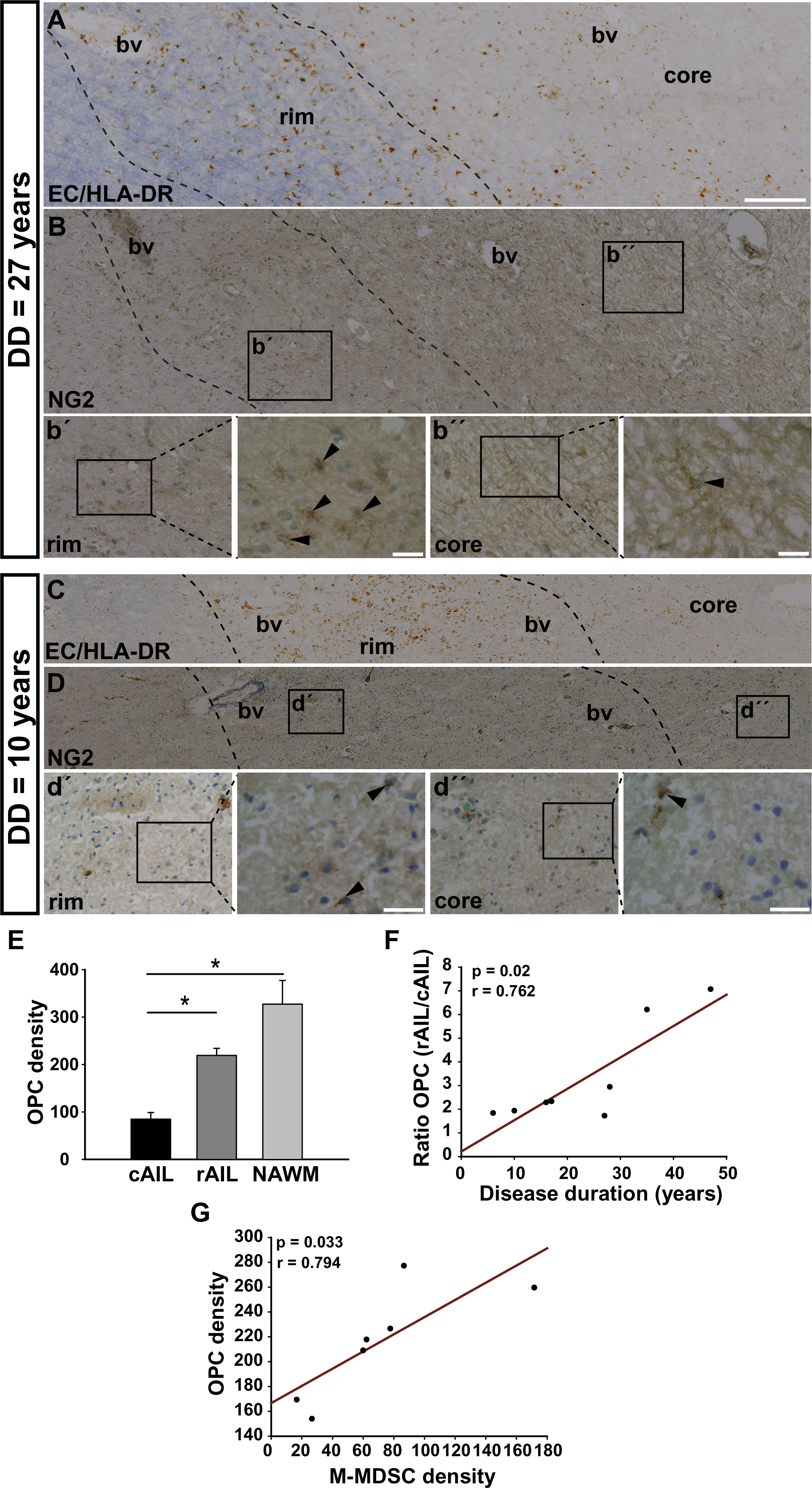
Disease duration affects the abundance of OPCs associated to mAIL in MS patients. **A-D:** Representative images of parallel sections of two mAIL from a MS patient with long disease duration (A-B) and a patient with short disease duration (C-D) showing myelin stained with EC (blue) and inflammatory cells immunostained with anti-HLA-DR (brown) in A, C, and OPCs identified as NG2^+^ cells in B, D. Magnification of the squared representative areas of the rim (b’, d’) and of the core (b’’, d’’) of the same patients showing OPCs pointed with arrowheads. **E:** Quantification of MS tissue samples showing a higher density of OPCs in the rAIL and in the NAWM compared to the cAIL (n = 8). **F:** Longer disease durations were associated with a higher ratio of OPC density in the rAIL/ OPC density in the cAIL (n = 8). **G:** The density of M-MDSCs in the rAIL was associated to a greater density of OPCs in the same area (n = 7). Scale bar: A, B = 200 μm; b’, b’’= 105 μm; C, D = 300 μm; d’, d’’ = 85 μm. bv, blood vessel; DD, disease duration. Rest of abbreviations as in the text.

**Table 1:**
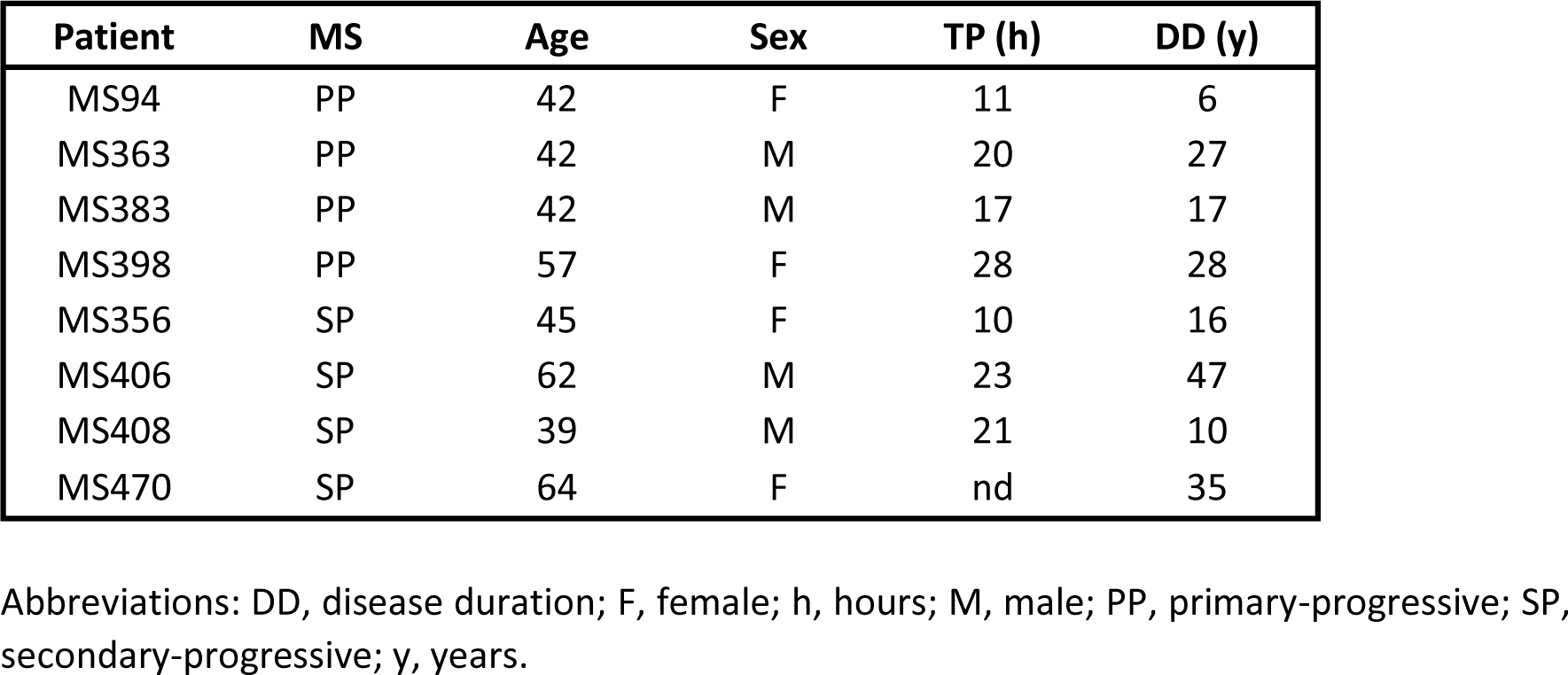
Demographic and clinical data of MS patients for the histopathological analysis.

| **Patient** | **MS** | **Age** | **Sex** | **TP (h)** | **DD (y)** |
| --- | --- | --- | --- | --- | --- |
| MS94 | PP | 42 | F | 11 | 6 |
| MS363 | PP | 42 | M | 20 | 27 |
| MS383 | PP | 42 | M | 17 | 17 |
| MS398 | PP | 57 | F | 28 | 28 |
| MS356 | SP | 45 | F | 10 | 16 |
| MS406 | SP | 62 | M | 23 | 47 |
| MS408 | SP | 39 | M | 21 | 10 |
| MS470 | SP | 64 | F | nd | 35 |

Abbreviations: DD, disease duration; F, female; h, hours; M, male; PP, primary-progressive; SP, secondary-progressive; y, years.

### The severity of the effector phase of EAE influences the abundance of M-MDSCs and OPCs in the demyelinated spinal cord

In order to investigate how disease severity affects the distribution of M-MDSCs and OPCs in the CNS of mice with EAE, we took advantage of the highly translatable individualized evaluation of this animal model [18,23]. In a first cohort of 12 mice, the severity index (SI) of EAE mice three days after disease onset ranged from 0.19 – 0.81, with a median of 0.64 (interquartile range-IQR 0.34 – 0.75). Animals were classified as mice with mild or severe EAE (n = 6 for each group), according to the median of the SI [19].

We wanted to evaluate the impact of the disease severity in the distribution of M-MDSCs and the level of OPC proliferation in the demyelinated spinal cord during the effector phase of EAE. For that, mice were administered BrdU for 3 consecutive days, starting at the specific disease onset of each animal, followed by the histopathological analysis of the spinal cord one day later (Figure 2A). Independently of disease severity, M-MDSCs (identified as Arg-I^+^ cells at this specific time point; [21,25,26]) were more abundant in the plaque of the lesions compared to the NAWM (Figure 2B, F, J), while OPCs (NG2^+^ cells) were predominantly found in the surrounding periplaque (Figure 2C, G, K). Moreover, proliferating OPCs (NG2^+^BrdU^+^ cells) were also more present in the periplaque of the lesions compared to the plaque irrespective of the disease severity. However, OPCs that incorporated BrdU during the effector phase were significantly more abundant in the periplaque than in the adjacent NAWM only in the case of mice with mild EAE (Figure 2D, E, H, I, L). Importantly, the severity of the effector phase drastically affected all of the cell populations mentioned above. M-MDSCs in the plaque as well as total and proliferating OPCs in the periplaque were more abundant in mice with a milder clinical course of EAE compared to those with severe EAE (Figure 2J-L). Consequently, we found that the milder clinical course according to the SI, the higher density of M-MDSCs in the plaque (Figure 3A), and of total and proliferating OPCs in the periplaque (Figure 3B-C). Therefore, these results indicate that the severity of the effector phase of the EAE clinical course negatively influences the abundance and proliferative state of OPCs within the demyelinated CNS. In the same sense, a milder disease severity positively impacts on the higher abundance of immunoregulatory M-MDSCs within the inflamed spinal cord lesions.

**Figure 2.**
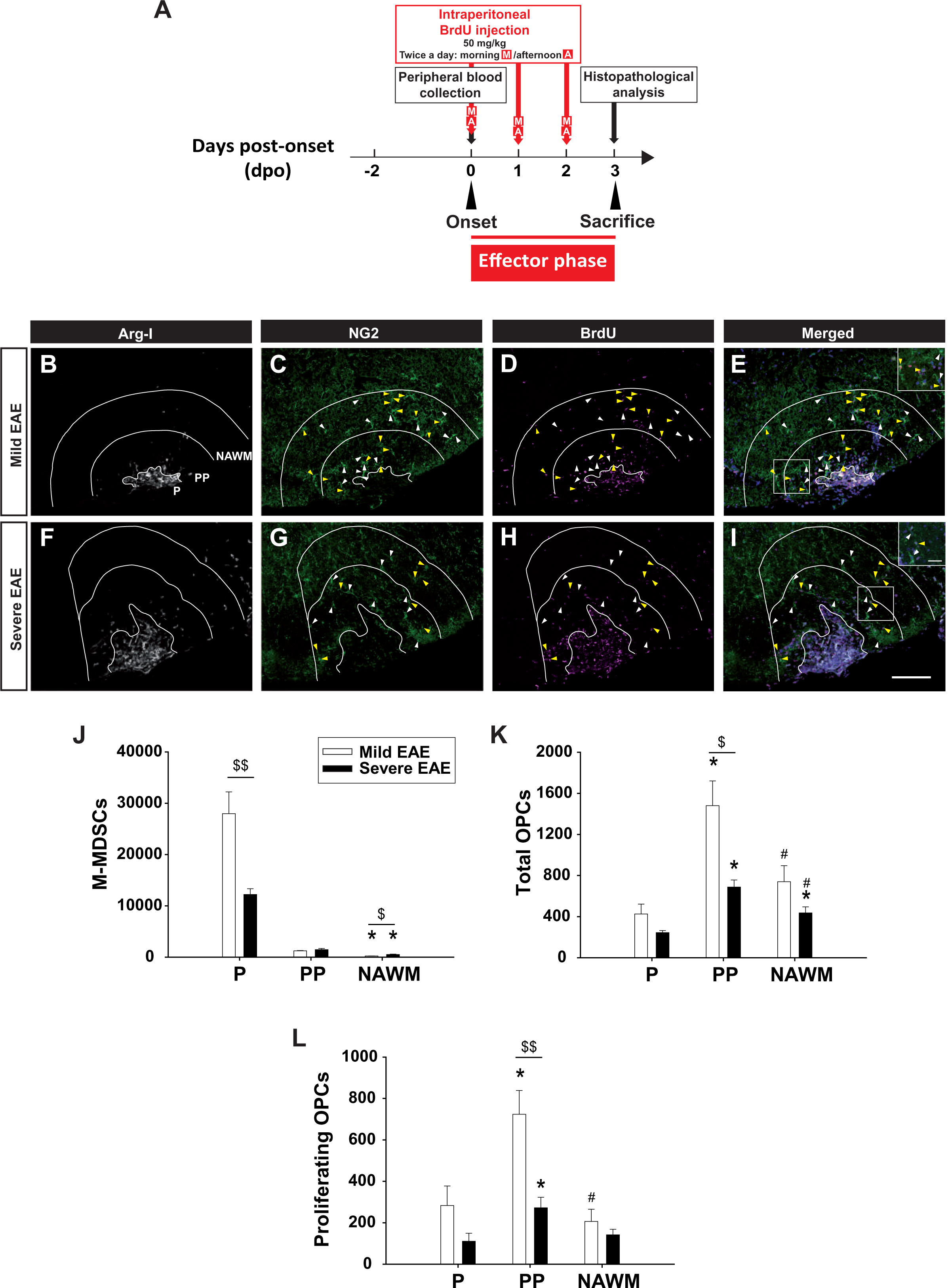
Disease severity impacts on M-MDSC presence and OPC generation and distribution within the demyelinated CNS at the peak of EAE. **A**: Diagram representing the experimental procedure. **B-I:** Representative confocal images showing the distribution of M-MDSCs (Arg-I^+^ cells, white), total OPCs (NG2^+^ cells, green, pointed with white arrows) and proliferating OPCs (BrdU^+^NG2^+^ cells, magenta, pointed with yellow arrows) in the plaque (P), periplaque (PP) and NAWM within the spinal cord of a mouse with a mild (B-E) or a severe (F-I) EAE. White lines indicate P, PP and NAWM borders. The ventral part is at the bottom and the lateral part is on the right. **J-L:** The density of M-MDSCs at the P (J), total OPCs (K), and proliferating OPCs (L) at the PP was higher in mice with a mild EAE compared to those with a severe disease course. ^$^p < 0.05; ^$$^p < 0.01, when comparing cell density between mice with mild *vs.* severe EAE; *p < 0.05, and ^#^p < 0.05 when comparing the cell density of an area with respect to the P or when comparing cell density of NAWM *vs.* the PP within a specific group of mice, respectively. Insets in E, I are higher magnifications of the white squares. Scale bar in B-I = 100 µm (20 µm in the insets).

**Figure 3.**
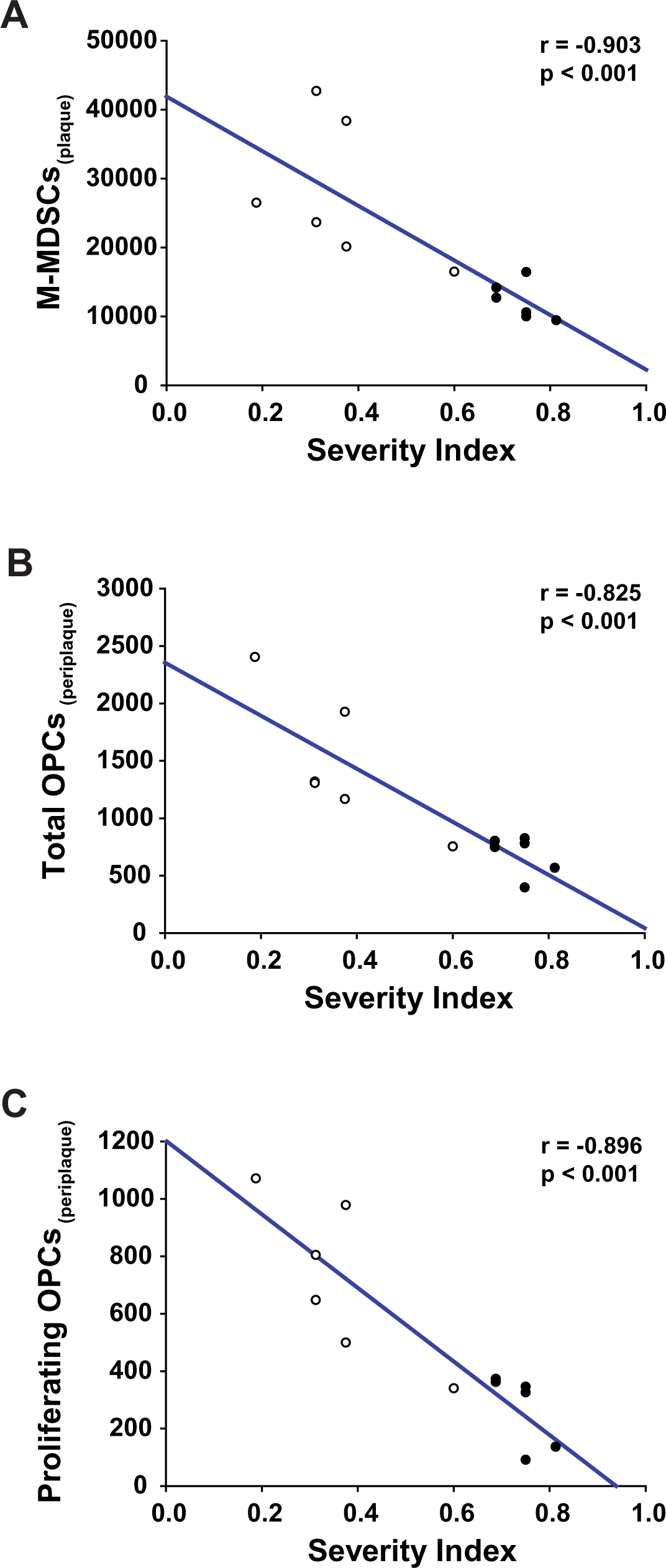
The higher severity of the effector phase of EAE is associated with a lower abundance of M-MDSCs and newly generated OPCs in the demyelinated lesions of the spinal cord. **A-C:** The higher severity of the effector phase of EAE was inversely correlated with the density of M-MDSCs in the plaque (P, A), the density of total OPCs (B) and proliferating OPCs (C) in the periplaque (PP). Spearman correlation test was carried out (n = 12). White circles represent mice with mild EAE (n = 6); black circles represent mice with severe EAE (n = 6).

### M-MDSC abundance is directly related to a higher OPC generation in the spinal cord of EAE mice

To explore the relationship between infiltrated M-MDSCs and both OPC abundance and proliferative activity, we correlated the density of M-MDSCs in the plaque, i.e. the area where they were more enriched, and the densities of OPCs and proliferating OPCs in the different areas of the demyelinated lesion. We observed a direct correlation between the density of M-MDSCs and the density of total and proliferating OPCs both in the plaque (Figure 4A, C) and in the periplaque (Figure 4B, D) but not in the NAWM (total OPCs: r = 0.483, p = 0.105; proliferating OPCs: r = 0.308, p = 0.317). Importantly, we also observed a direct correlation between M-MDSCs abundance and a gradient of proliferating OPCs between the periplaque and the adjacent NAWM (Figure 4E). Therefore, the presence of M-MDSCs in the plaque is directly related to a higher abundance and proliferative activity of OPCs in the closely related areas at the end of the effector phase of EAE.

**Figure 4.**
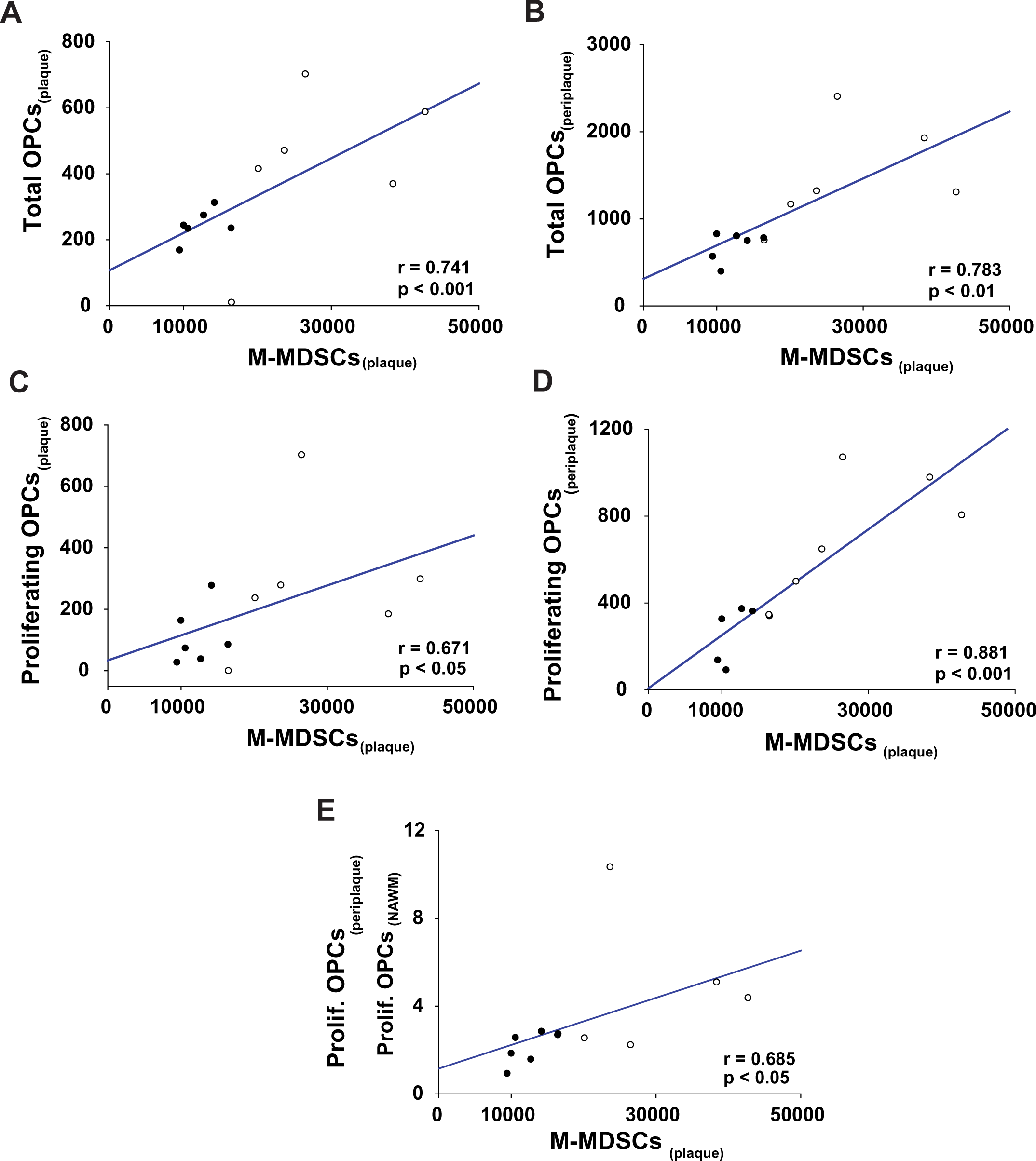
The densities of M-MDSCs and OPCs strongly interrelate in the spinal cord of EAE mice. **A, B:** The abundance of M-MDSCs in the plaque (P) was associated with the density of OPCs in the plaque (P, A) and in the periplaque (PP, B). **C, D:** The higher density of M-MDSCs in the P, the higher density of proliferating OPCs in the P (C) and in the PP (D). **E:** M-MDSC distribution also associated with a gradient of proliferating OPCs between the PP and the adjacent NAWM. Spearman correlation test was carried out (n = 12). White circles represent mice with mild EAE (n = 6); black circles represent mice with severe EAE (n = 6).

We wanted to know the ability of circulating M-MDSCs as biomarkers of OPC density and proliferative activity within the inflamed spinal cord of EAE mice. First, we confirmed that the higher abundance of M-MDSCs in peripheral blood at the onset of EAE, the milder disease courses according to the SI (r = -0.775, p< 0.01; [23]). Interestingly, we found that the more abundance of M-MDSCs in peripheral blood at disease onset, the more density of these cells in the plaque of the demyelinated lesions at the peak of the disease (r = 0.762, p < 0.01). With respect to OPCs, the circulating M-MDSC load at disease onset tended to correlate with the higher abundance of total and proliferating OPCs both in the plaque (r = 0.497, p = 0.09, for total OPCs; and r = 0.538, p = 0.07, for proliferating OPCs) and in the periplaque of the lesions (r = 0.531, p = 0.07, for total OPCs), being nearly significant in the case of proliferating OPCs (r = 0.566, p = 0.05). In sum, central and peripheral M-MDSCs are positively associated to a higher content of total and newly generated OPCs in spinal cord lesions at the end of the pro-inflammatory phase of EAE.

### The density of OPCs in demyelinated EAE lesions at the end of the recovery phase is independent of disease severity

In a next step we wondered whether the impact of EAE severity on the distribution of OPCs generated during the effector phase or at the beginning of the recovery phase of EAE might still be observable in demyelinating spinal cord lesions once each animal reached its maximal clinical recovery. To this aim, a second cohort of mice with EAE (n = 12) was again distributed in two different groups according to the median of the SI [ranged from 0.50 – 0.83, with a median of 0.69 (IQR = 0.60 – 0.75)]. Following the same criteria than for the first animal cohort, animals with a SI < median were classified as mice with mild EAE (n = 5), while those with a SI > median were classified as mice with severe EAE (n = 7). In this new cohort, we traced cell division during both the whole effector and the beginning of the recovery phases of EAE by injecting BrdU from the onset until the peak of the disease, and then EdU in the same way for 3 consecutive days (Figure 5A). Once the animals reached their maximal functional recovery, we quantified the density of total OPCs (NG2^+^ cells) and the density and percentage of proliferating OPCs that came both from the effector phase (NG2^+^BrdU^+^cells) or from the early recovery phase (NG2^+^EdU^+^ cells; Figure 5B-K). Unlike that observed at the peak of EAE, we did not observe any difference in OPC distribution between regions in the lesions of both animals with mild and severe disease courses except for a higher abundance of total OPCs in the NAWM compared to the plaque in mice with severe EAE (Figure 5L). Moreover, the densities of total OPCs and proliferating OPCs were not different between mild and severe EAE mice (Figure 5L-M). Consequently, the percentage of proliferating OPCs and the ratio between proliferating and non-proliferating OPCs were also similar in animals with high or low severity of the clinical course (Figure 5N-O). These data show that the abundance of OPCs in the demyelinated spinal cord lesions once each animal has reached the maximal clinical recovery do not reflect the preceding severity of the clinical course of EAE.

**Figure 5.**
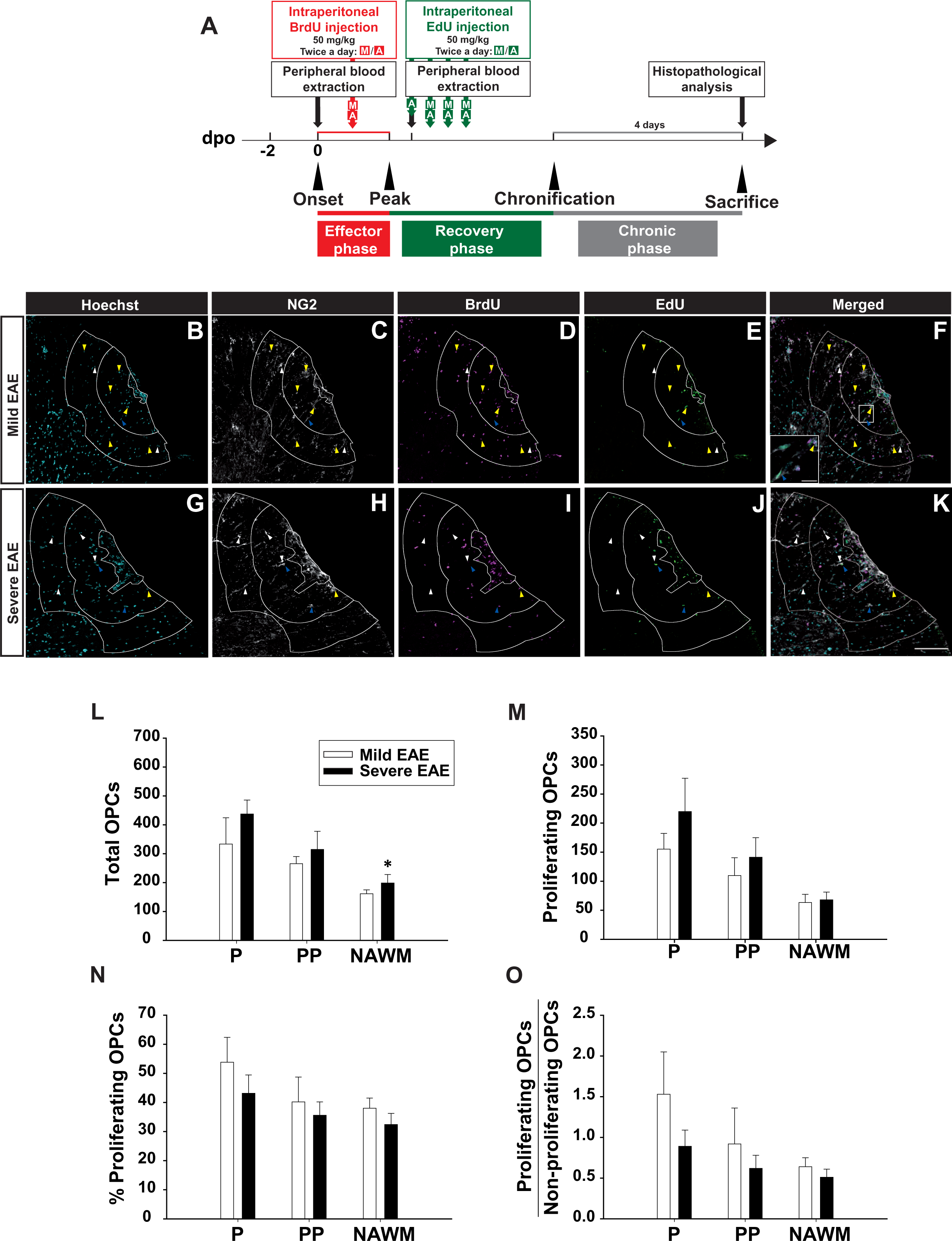
Disease severity does not mirror the distribution of OPCs present in the demyelinated spinal cord at the end of the recovery phase. **A:** Diagram representing the experimental procedure. **B-K:** Representative confocal images showing the distribution of total OPCs (NG2^+^ cells, white, white arrowheads), OPCs originated during the effector phase of EAE (NG2^+^BrdU^+^ cells, magenta, yellow arrowheads) and OPCs generated during the recovery phase of EAE (NG2^+^EdU^+^ cells, green, blue arrowheads), in the plaque (P), periplaque (PP) and NAWM within the spinal cord of mice that experienced a mild (B-F) or a severe (G-K) EAE. Cell nuclei was visualized with Hoechst (B, G). White lines indicate P, PP and NAWM borders. Ventral is on the right and lateral is on the top. **L-O:** Quantification of total (L) and proliferating OPCs (M) densities did not reveal any difference between EAE groups. Mice with severe EAE exhibited a slightly greater abundance of OPCs in the NAWM compared to the P (L). **N, O:** Neither the percentage of proliferating OPCs (NG2^+^BrdU^+^ or NG2^+^EdU^+^ cells, N), nor the ratio between proliferating and non-proliferating OPCs (O) were affected by disease severity. *p < 0.05, when comparing the cell density of an area with respect to the P within a specific group of mice. Inset in F is a higher magnification of the white square, showing NG2^+^BrdU^+^ cells (yellow arrowheads) and NG2^+^EdU^+^ cells (blue arrowheads). Scale bar in B-K = 100 μm (15 μm in the inset).

### Disease severity influences mature oligodendrocyte generation in the spinal cord of EAE mice

We wanted to assess the influence of disease severity in OL generation in the demyelinated spinal cord of EAE mice. Once the animals had reached their maximal functional recovery, we quantified the density of total OLs (CC1^+^Olig2^+^ cells), newly generated OLs during the whole effector phase (CC1^+^Olig2^+^BrdU^+^) or during the early recovery phase (CC1^+^Olig2^+^EdU^+^ cells), and surviving OLs (CC1^+^Olig2^+^ BrdU^-^EdU^-^ cells; Figure 6A-J). We did not find any difference in the cellular densities of total or surviving OLs within the periplaque of animals with mild or severe EAE (total OLs: mild EAE = 430.11 ± 83.02, severe EAE = 278.34 ± 40.49, p = 0.102; surviving OLs: mild EAE = 299.13 ± 59.41, severe EAE = 219.00 ± 29.39, p = 0.215). Interestingly, we found an increased production of OLs in the periplaque of mice with milder disease courses irrespective of the clinical phase of origin (Figure 6K, L). In general, the percentage of newly generated OLs with respect to the total OL population was higher in the periplaque of mice with mild EAE than those with severe EAE (Figure 6M). Of note, the ratio between newly generated OLs and surviving OLs was also higher in the periplaque of the group of EAE animals with milder disease courses (Figure 6N). Overall, these results indicate that disease severity importantly affects OL generation in the periplaque of spinal cord demyelinated lesions of EAE mice.

**Figure 6.**
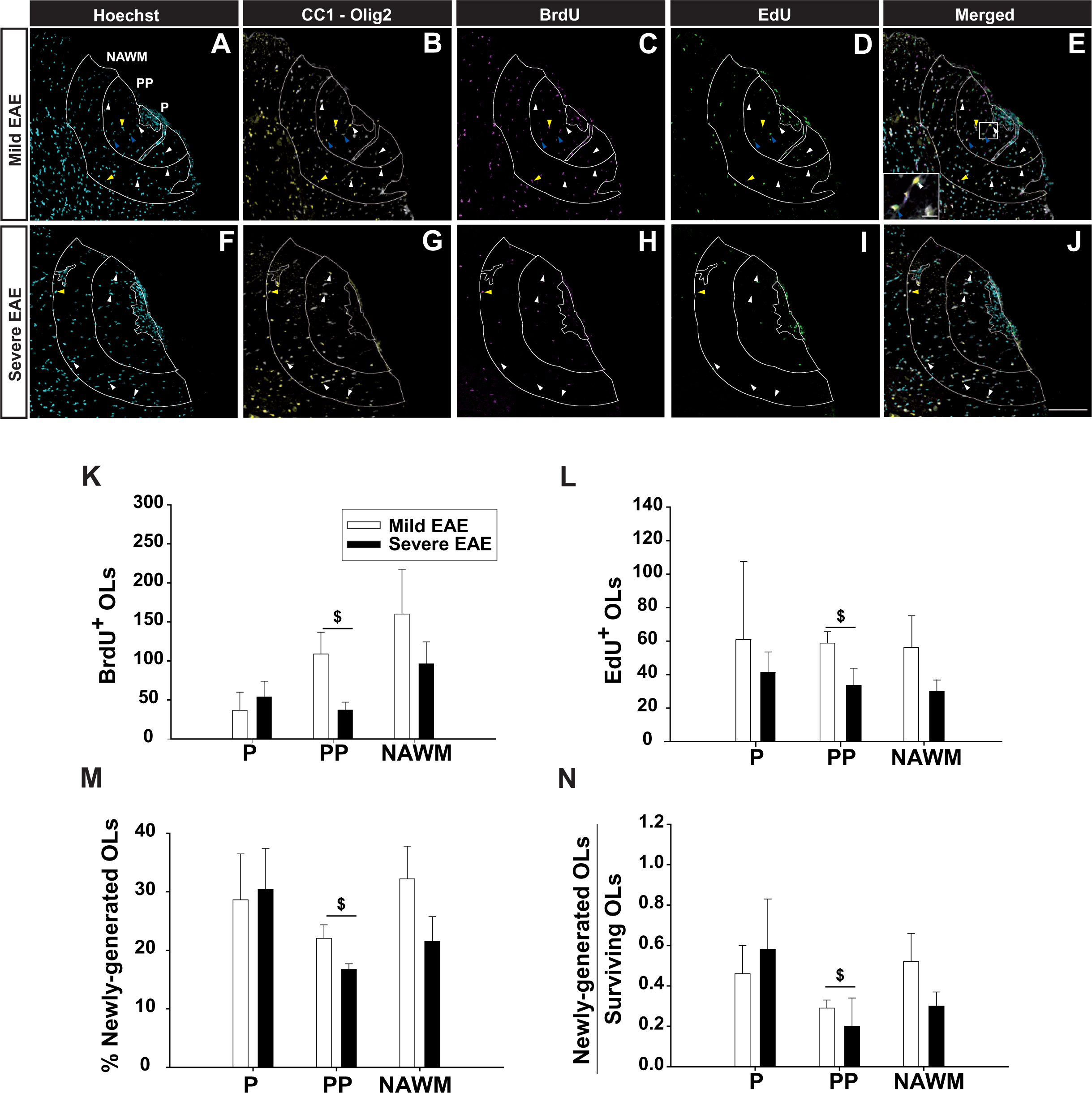
The severity of EAE affects the distribution of newly generated oligodendrocytes in the demyelinated spinal cord after the recovery from the inflammatory insult. **A-J:** Representative confocal images showing the distribution of mature OLs (CC1^+^Olig2^+^ cells, white cytoplasm with yellow nucleus, white arrowheads), OLs originated during the effector phase of EAE (CC1^+^Olig2^+^BrdU^+^ cells, white cytoplasm with magenta nucleus, yellow arrowheads) and OLs generated during the early recovery phase of EAE (CC1^+^Olig2^+^EdU^+^ cells, white cytoplasm with green nucleus, blue arrowheads), in the plaque (P), periplaque (PP) and NAWM within the spinal cord of mice with a mild (A-E) and a severe (F-J) EAE. Cell nuclei was visualized with Hoechst (A, F). White lines indicate P, PP and NAWM borders. Ventral is on the right and lateral is on the top. **K-N:** The density of BrdU^+^ (K) and EdU^+^ newly originated Ols (L) was higher in the PP of mice with mild EAE compared to those with a severe disease. **M:** Mice with a mild EAE presented a higher percentage of newly generated OLs (CC1^+^Olig2^+^BrdU^+^ or CC1^+^Olig2^+^EdU^+^ cells) in the PP than mice that experienced a severe EAE. **N:** The ratio between newly generated and surviving OLs was also higher in the PP of mice with mild vs. severe EAE. ^$^p < 0.05. Inset in E is a higher magnification of the white square. Scale bar: A-J = 100 μm (15μm in the inset).

### Circulating M-MDSC load is indicative of the future production of mature OLs in spinal cord demyelinated lesions

We previously reported that peripheral blood M-MDSCs at the onset of the EAE is associated with milder disease courses and less demyelination and axonal damage in the spinal cord of EAE mice [23]. Here, we corroborated the inverse correlation between circulating M-MDSC load in blood at disease onset and the future SI (r = -0.582, p < 0.05). Moreover, we found that the higher abundance of M-MDSCs in peripheral blood at onset, the higher recovery index (r = 0.633; p < 0.05), which is indicative of a faster reduction of EAE symptoms. Finally, we confirmed that the higher load of circulating M-MDSCs at the peak of the disease, the greater and faster recovery of EAE mice (percentage of recovered score: r = 0.664, p < 0.05; recovery index: r = 0.692, p < 0.05). These results show that the abundance of circulating M-MDSCs both at the onset and at the peak of the disease is indicative of a better symptom recovery.

We next explored the predictive ability of circulating M-MDSCs on the presence and generation of OPCs and OLs in the plaque and the periplaque of lesions at the end of the recovery phase. Regarding OPCs, we did not find any correlation between the percentage of circulating M-MDSCs at the onset or the peak of EAE with the density of total or proliferating OPCs both in the plaque and in the periplaque of the lesions (data not shown). On the contrary, the proportion of circulating M-MDSCs at EAE onset directly correlated with a plaque enriched in newly generated OLs at the end of the recovery phase (Figure 7A). Interestingly, the abundance of circulating M-MDSCs at the peak of EAE was directly associated with the percentage of newly generated OLs (Figure 7B) as well as with the enrichment of newly generated OLs in the periplaque (Figure 7C). These data indicate that the proportion of M-MDSCs in peripheral blood of mice with EAE can give information about the future greater ability of the CNS to produce mature OLs associated to demyelinated areas.

**Figure 7.**
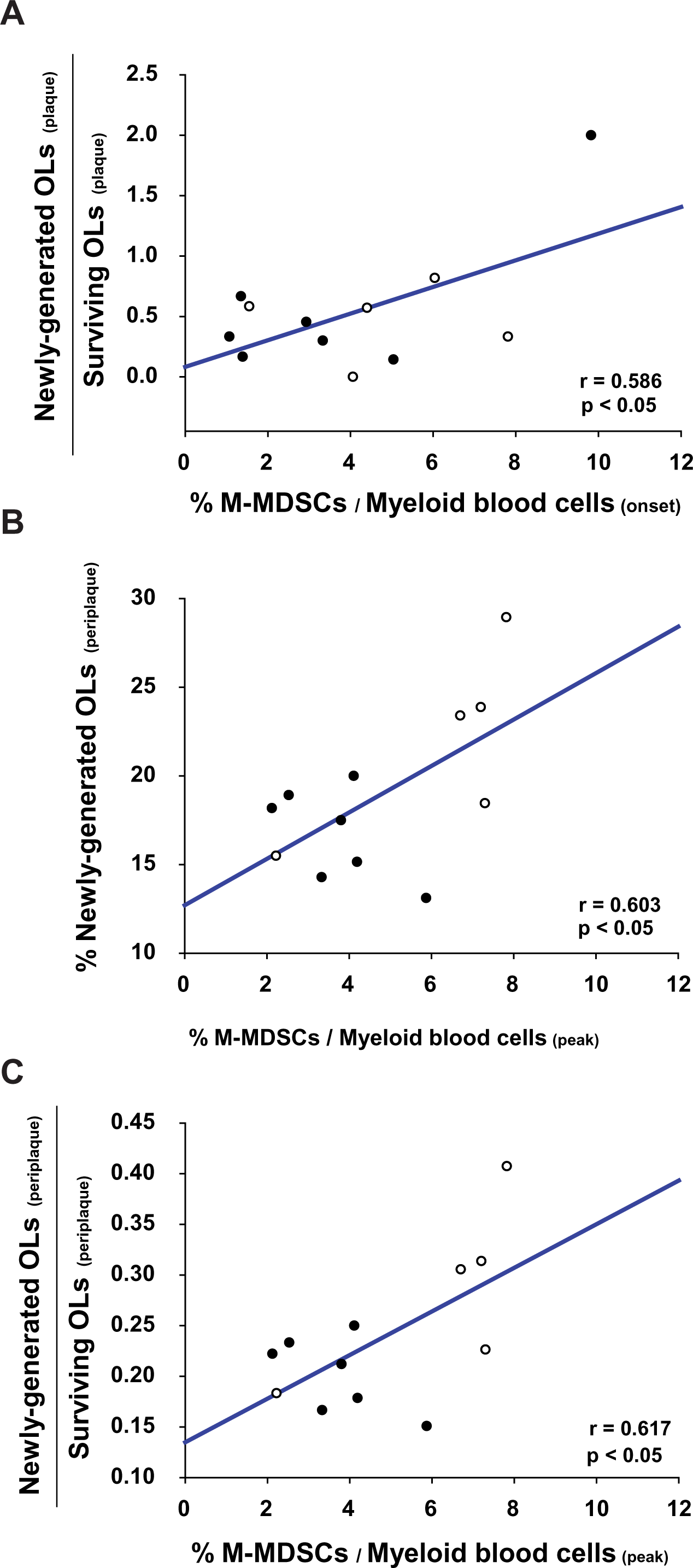
Circulating M-MDSC load is indicative of a CNS prone to produce new OLs during the disease course of EAE. **A:** The abundance of circulating M-MDSCs at EAE onset is directly associated with a higher presence of newly generated OLs in the plaque of the lesions at the end of the recovery phase. **B:** The higher proportion of M-MDSCs in peripheral blood at the peak of EAE, the higher percentage of newly generated OLs in the periplaque of the lesions. **C:** The abundance of circulating M-MDSCs at the peak of EAE directly correlates with a greater enrichment of newly generated OLs in the periplaque of the lesions. Pearson correlation test was carried out (n = 12). White circles represent mice with a mild EAE (n = 5); black circles represent mice with a severe EAE (n = 7).

## DISCUSSION

Inefficient remyelination in MS leads to neurodegeneration, which is closely related to the permanent disability observed in MS patients [6,11]. To date, the majority of the available treatments are indicated for reducing disease activity (relapses) or the associated symptoms but do not tackle neurodegeneration. Hence, there is an urgent need for the search of novel compounds that enhance myelin regeneration and prevent the associated axonal degeneration [13]. Despite the large variety of pro-remyelinating candidate drugs, only a few have reached the Phase 2 clinical trials. However, those trials completed in RRMS patients have shown little or no clinical improvements [27–29]. This could be due to interindividual variability in the ability of oligodendrocyte generation after demyelination of each MS patient. Differentiation of OPCs into OLs seems to be a pre-requisite for remyelination [30–33] and recent studies point newly generated OLs as more efficient remyelinating cells than those that survive demyelination [34,35]. Thus, it is a priority to find suitable biomarkers that allow clinicians to better classify patients according to their ability to produce new OPCs able to differentiate into myelinating OLs.

In the present work, we demonstrate in MS tissue that longer disease durations are closely related to a higher relative abundance of OPCs in the rim *versus* the core of mAIL. Interestingly, a higher abundance of M-MDSCs is directly related to a greater density of OPCs in the same lesions. Concurrently, we show that the disease severity of the EAE affects the distribution and proliferative activity of oligodendroglial cells in spinal cord demyelinating lesions. We observe that milder disease courses in EAE are associated with a higher abundance of new OPCs in the periplaque of the lesions at the end of the effector phase and also with a greater presence of new OLs in the same area once each animal complete their maximal symptom recovery. We find that the higher the density of infiltrated M-MDSCs within the plaque of the lesions, the greater presence of newly generated OPCs in the periplaque at the peak of the EAE clinical course. Finally, we show the association between circulating M-MDSC abundance and the enrichment of newly generated OLs in the lesions after EAE recovery.

We observe a higher ratio of OPC density in the rAIL/ OPC density in the cAIL, which means that milder forms of the disease are associated with a higher presence of OPCs in the rim of the lesions, the same result that we find in the EAE model. In patients with progressive MS, a greater proportion of mAIL is associated to a shorter disease duration together with a shorter time to reach an expanded disability status scale (EDSS) of 6 or 8 [36]. Therefore, our data suggest that disease severity negatively influences OPC abundance in those lesions associated to more aggressive MS courses. Moreover, we observe that the higher density of M-MDSCs, the higher density of OPCs in the rAIL, which is in accordance with our previous observations in the EAE model, where the presence of M-MDSCs in the plaque of the lesions directly correlated with the density of OPCs in the surrounding periplaque [22]. M-MDSCs have been recently found in highly inflammatory areas of progressive MS patients, including the rAIL. Interestingly, in primary-progressive MS patients, a lower content of M-MDSCs together with a more evidenced pro-inflammatory environment is associated with a more severe disease course [23]. Therefore, our data suggest that M-MDSCs may have a relevant role for OPC biology in the context of MS demyelinating lesions.

EAE mice present a biological variability that resembles the reality of MS patients in relation to the heterogeneity of disease severity [19]. Our group has shown that the individualized follow-up of the animals constitutes a good strategy for the search of biomarkers in this MS model, a scientific formula that helps to close the gap between preclinical and clinical research [18,23]. In the present work, the individualized monitoring of the EAE mice allow us to demonstrate that EAE severity influences OPC distribution and self-renewal at the peak but not at the recovery phase of the disease, while OPC differentiation is highly detected in animals with a mild EAE once they reached their maximal symptom recovery. Namely, at the peak of EAE, mice with a mild disease course present a greater density of total and proliferating OPCs in the periplaque of the spinal cord lesions originated during the effector phase of the disease when pro-inflammatory myeloid cells are predominant [37,38]. This is consistent with the fact that pro-inflammatory molecules released by microglia/macrophages, but also by OPCs themselves, stimulate remyelination by enhancing OPC activation and proliferation [39–42]. In addition, we find a direct association between the presence of M-MDSCs at the plaque and the abundance of proliferating OPCs in the periplaque, which is in line with our previous work that describe M-MDSCs as promoters of OPC proliferation *in vitro* [22]. Furthermore, we observe a higher percentage of newly generated OLs in the periplaque from mice with a mild course of EAE originated both during the effector phase or the early immunomodulatory phase preceding symptom recovery. The latter examined period marks the switch of monocyte-derived macrophages from a pro-inflammatory to an anti-inflammatory profile [37,38], a cellular state with a proved ability to induce the differentiation of OPCs [43].

We previously observed that the severity of EAE is related to several histopathological hallmarks, as mice with severe disease courses present more lymphocyte infiltration, demyelination and axonal damage [19]. Thus, although a moderate inflammation is crucial to drive myelin-associated regenerative processes [42,44], a persistent pro-inflammatory environment [15,45] together with a poor presence of cells with anti-inflammatory properties [43,46,47] could contribute to OPC differentiation arrest, this being the case in the active lesions and in the rim of mAIL from MS patients [48]. Remyelination is almost absent at the peak of EAE while it is observed at the chronic stage, which indicates that the regeneration of myelin fails in the presence of an exacerbated inflammation [15], something that may be reversed by pharmacologically modulated pro-inflammatory cells [33]. However, an anti-inflammatory milieu able to reduce EAE severity and potentiate the immunosuppressive activity of regulatory cells as M-MDSCs, could prompt myelin regeneration, as these cells are associated with increased T cell apoptosis [19] and modulate OPC proliferation and differentiation [22]. In addition to the influence of the inflammatory/anti-inflammatory environment, it has been described a clear OPC heterogeneity dependent on origin, age and local environment both in homeostatic or pathological conditions [49–51]. Therefore, we cannot rule out that the existence of an interindividual variability in the density of OPCs and/or OLs previous to the demyelinating insult may affect, at least in part, the endogenous ability of each mouse to produce new oligodendroglial cells.

Notably, we find a positive relationship between circulating M-MDSCs at the onset and peak of EAE and the enrichment of newly generated OLs in the plaque and periplaque of the lesions, respectively, once each animal reached the maximal recovery. Circulating M-MDSCs at disease onset have been shown to be an excellent biomarker of milder disease courses and less demyelination and axonal damage in the spinal cord at the end of the effector phase of EAE [23]. Moreover, it was shown that the greater abundance of circulating M-MDSCs at the peak of EAE, the better symptom recovery in the same MS model [23]. Thus, circulating M-MDSCs in EAE point to be not only a promising biomarker for predicting milder disease courses, but also for indicating a CNS prone to produce new OLs, whose interrelationship is highlighted in the present work. Despite the substantial advances made in myelin neuroimaging techniques using magnetic resonance imaging (MRI) and positron emission tomography (PET; [52]), they are still not fully validated histopathologically and have inherent characteristics (technical, budgetary, image post-processing) that limit their utility for clinical routine. Moreover, the observations made with these technical approaches cannot be easily extrapolated to clinical studies due to the absence of biochemical or imaging biomarkers for recently differentiated OLs and new myelin formation that allow clinicians to easily recognize myelin regeneration. In this sense, our study may represent a key starting point to find this kind of biomarkers which should be corroborated in future clinical studies associating M-MDSC abundance with myelin restoration.

## CONCLUSIONS

Here we show for the first time that disease severity is associated with the abundance of OPCs present in mAIL from MS patients and with the generation of OLs in the MS animal model EAE. Furthermore, our work shows a relationship between circulating M-MDSCs and the future OL generation in demyelinated spinal cord lesions found at the end of the clinical symptom recovery. This study sets the basis for the preclinical search for new biomarkers aimed at identifying a CNS prone to the generation of new OPCs and OLs that may contribute to restore the damaged myelin, something with relevant implications for the future development of pro-myelinating treatments.

## METHODS

### Human tissue and MS lesion classification

*Post-mortem* snap frozen cortical tissue blocks containing mAIL from 8 MS patients were analyzed (supplied by the UK MS Tissue Bank; Table 1). mAIL were classified according to demyelination and cellular distribution as previously described [36,53,54]: demyelinated areas with a hypocellular cAIL surrounded by a rAIL enriched with human-leukocyte antigen (HLA)- DR positive cells. Moderate T cell infiltration was observed in the center of mAILs.

### Immunohistochemistry and eriochrome cyanine for myelin staining in human tissue

Cryosections (10 µm, Leica) from MS tissue were air-dried for 45 min at room temperature (RT) and fixed in 4 % paraformaldehyde (PFA; Sigma Aldrich) in 0.1 M Phosphate Buffer, pH 7.4 (PB) for 1 h. Immunohistochemistry (IHC) for HLA-DR and OPCs was carried out by incubating with anti-human HLA-DR (1:200; Agilent, M0746, TAL.1B5 clone) and anti-human NG2 (1:25; clone 9.2.27, BD Bioscience) individually in parallel tissue slides. Detection of NG2 was visualized using TSA signal amplification system (TSA Plus biotin kit, AkoyaBio) and the appropriate biotinylated secondary antibodies (1:200; Vector Labs) were used. The IHC reaction was developed using the Vectastain Elite ABC reagent (Vector Labs) and the peroxidase reaction product was visualized with 0.05 % 3, 3’-diaminobenzidine (DAB, Sigma-Aldrich) and 0.003 % H_2_O_2_ in 0.1 M Tris-HCl, pH 7.6. The reaction was monitored under the microscope and terminated by rinsing the slides with PB. After NG2 IHC, cell nuclei were counterstained with Agilent Giemsa Stain Kit (AR164, Artisan). To visualize myelin after HLA-DR IHC, eriochrome cyanine (EC) staining was carried out as described elsewhere [23]. In all cases, the sections were dehydrated and mounted with Entellan mounting medium.

M-MDSCs were identified in parallel tissue slices from the same patients by immunofluorescence with a combination of three markers to distinguish these cells from other myeloid cells such as neutrophils or inflammatory macrophages as described previously [23]: the monocytic marker CD14 (always expressed by M-MDSCs, 1:25, R&D, BAF383), the granulocytic protein CD15 (always absent in M-MDSCs given that it is observed in both mature neutrophils and the polymorphonuclear subset of MDSCs, 1:25, Agilent ISO62, carb3 clone) and the specific molecule of antigen presenting-cells HLA-DR (low or absent in M-MDSCs; 1:100, Agilent M076, TAL.1B5 clone). After incubating the tissue with the primary antibodies, the appropriate Alexa fluorescent-tagged secondary antibodies were used (1:1000, Invitrogen) and cell nuclei were stained with Hoechst 33342 (10 µg/mL, Sigma-Aldrich).

### OPC and M-MDSC quantification in mAIL

By using HLA-DR IHQ and EC, rAIL was defined as the region between the border of the demyelinated core and the adjacent non-demyelinated white matter enriched with HLA-DR^+^ cells. We also defined the NAWM as the non-lesional white matter one high power field (HPF) 40X (0.0485 mm^2^ per HPF) away from the end of the rAIL [55]. Only those mAIL where the 3 regions were clearly identified (cAIL, rAIL, and NAWM) were quantified. We randomly selected 5 HPF per region in the slide with NG2 immunostaining. Those elements with Giemsa positive nuclei surrounded by NG2 labelling were considered as OPC (NG2^+^ cells). The average of OPCs per region was determined and the density of the OPCs/mm^2^ of the HPF was calculated. HPF at 40X magnification of the three different regions of mAILs were captured in a BX61 microscope (Olympus) connected to a MBF CX9000 color camera. Given that there is not a specific marker for the identification of M-MDSCs in the tissue, cells with CD14^+^HLA-DR^-/lo^CD15^-^ phenotype were explored for the quantification of M-MDSC density in the rAIL as described previously [23].

### Induction of EAE

Chronic progressive EAE was induced in female 6- to 7-week-old C57/BL6 mice (Janvier Labs, France) by immunization with 200 µL of a Myelin Oligodendrocyte Glycoprotein (MOG) solution (200 µg, MOG_35-55_ peptide: Genscript, New Jersey, USA) emulsified in complete Freund’s Adjuvant (CFA) containing 4 mg/mL of heat inactivated *Mycobacterium tuberculosis* (BD Biosciences, Franklin Lakes, New Jersey, USA). Immunized mice were administered Pertussis toxin intravenously through the tail vein (250 ng/mouse: Sigma-Aldrich, St. Louis, MO, USA) in a final volume of 100 µL on the day of immunization and 48 h later. EAE was scored clinically on a daily basis in a double-blind manner as follows: 0, no detectable signs of EAE; 0.5, half limp tail; 1, whole limp tail; 1.5, hind limb inhibition (unsteady gait and/or poor hind leg grip); 2, hind limb weakness; 2.5, bilateral partial hind limb paralysis or unilateral full hind limb paralysis; 3, complete bilateral hind limb paralysis (paraplegia); 3.5, partial forelimb paralysis; 4, tetraplegia; 4.5, moribund; and 5, death. In accordance with ethical regulations, humane end-point criteria were applied when an animal reached a clinical score ≥ 4, when clinical score ≥ 3 was reached for more than 48 h, or when self-mutilation was evident, persistent retention of urine, 35 % weight loss and signs of stress or pain for more than 48 h, even if the EAE score was < 3. All animal manipulations were approved by the institutional ethical committee (*Comité Ético de Experimentación Animal del Hospital Nacional de Parapléjicos*), and all experiments were carried out in compliance with the European guidelines for animal research (European Council Directive 2010/63/EU/, 90/219/EEC, Regulation No. 1964/2003), and with the Spanish National and Regional Guidelines for Animal Experimentation (RD 53/2013 and 178/2004, Ley 32/2007 and 9/2003, Decreto 320/2010).

### Evaluation of EAE clinical signs

The clinical parameters analyzed were defined as: (i) the day of onset, i.e. the first day mice had a clinical score ≥ 0.5; (ii) clinical score at onset; (iii) the maximum clinical score (“peak”); (iv) the day when animals reached the peak; (v) the SI, quantified as the ratio between the maximum clinical score and the days elapsed from onset to peak [19]; (vi) the recovered score (the maximum clinical score – the final score); (vii) the percentage of recovery [(the recovered score * 100) / the maximum clinical score]; (viii) the recovery index [(the maximum clinical score – the final score) / the days elapsed from peak to chronification]; and (ix) the final score, the clinical score at the end of the follow-up.

### BrdU/EdU incorporation and follow-up in EAE mice

At the individualized onset of the clinical signs of each animal BrdU (50 mg/kg in sterile 1x Phosphate Buffered Saline - PBS) was intraperitoneally (ip) injected twice a day during 3 consecutive days in the first animal cohort (n = 12) or until the peak of the clinical course in the second animal cohort (n = 12). In this last cohort, once the animals reached the peak, after 24 h of wash out, EdU was administered ip (50 mg/kg in PBS) twice a day during 3 consecutive days. After EdU administration, animals were sacrificed once they reached their maximal functional recovery (chronification: when they repeated the same score at least for 4 consecutive days).

### Flow cytometry analysis of peripheral blood

Blood was collected from the submandibular vein of isofluorane-anesthetized mice at disease onset as well as at the peak of the clinical course in 2 % EDTA tubes. Erythrocytes were then lysed in a 15-mL tube with 2.5 mL ACK lysis buffer: 8.29 g/L NH4Cl; 1 g/L KHCO_3_; 1 mM EDTA in distilled H_2_O at pH 7.4 (Panreac). Subsequently, the lysis reaction was stopped by adding PBS and the blood cells were recovered by centrifugation at 210 g for 5 min at RT. The Fc cell receptors were then blocked for 10 min at 4 °C with and anti-CD16/anti-CD32 antibodies (10 µg/mL: BD Biosciences 553142) and the cells were resuspended in 25 µL of staining buffer: PBS supplemented with 10 % heat-inactivated fetal bovine serum (FBS: Capricorn; 25 mM), 2.5 % HEPES buffer (Gibco), 2 % Penicillin/Streptomycin (P/S: Gibco) and 0.2 % EDTA (Sigma; 0.5 M). The cells were further labeled for 30 min at 4 °C in the dark with the following fluorochrome-conjugated monoclonal antibodies diluted in 25 µL of staining buffer: rat anti-mouse Ly-6C-FITC (10 µg/mL, AL-21 clone), rat anti-mouse Ly-6G-PE (4 µg/ml, 1A8 clone), rat anti-mouse CD11b-PerCP-Cy5.5 (4 µg/ml, M1/70 clone: all from BD Biosciences), rat anti-mouse MHC-II-PE-Cy7 (4 µg/ml, M5/114.15.2 clone), hamster anti-mouse CD11c-APC (4 µg/ml, N418 clone), and rat anti-mouse F4/80-eFluor450 (4 µg/ml, BM8 clone) to stain myeloid cells (eBioscience-Thermo Fisher Scientific). The blood cells were then rinsed with staining buffer, recovered by centrifugation at 210 g for 5 min and fixed with 4 % PFA in PB for 10 min at RT. Analysis was performed in a FACS Canto II cytometer (BD Biosciences) at the Flow Cytometry Core Facility of the *Hospital Nacional de Parapléjicos*. The data obtained were analyzed by using the FlowJo 10.6.2 software (Tree Star Inc.).

### Tissue sampling and immunohistochemistry for murine tissue

Animals were perfused transcardially with 2 % PFA and their spinal cords were dissected out and post-fixed for 4 h at RT in the same fixative. After immersion in 30% (w/v) sucrose in PB for 12 h, seriated coronal cryostat sections (20 µm thick: Leica, Nussloch) were thaw-mounted on Superfrost® Plus slides and stored at -20°C until use. Spinal cord sections were air-dried for 1 h at RT. After several rinses with PB, the sections were preincubated for 1 h at RT in blocking buffer: 5% normal donkey serum (NDS, Vector) and 0.2 % Triton X-100 (Merck) diluted in PBS. IHC was carried out by incubating the sections overnight at 4 °C with the following primary antibodies diluted in blocking buffer: rabbit anti-NG2 (1:200; AB5320, Millipore), mouse anti-CC1 (1:200; OP80, Calbiochem) and rabbit anti-Olig2 (1:200; AB9610, Millipore). The sections were kept for 30 min at RT and, after rinsing, antibody binding was detected with the corresponding fluorescent secondary antibodies in incubation buffer for 1 h at RT (1:1,000, Invitrogen) and the sections were fixed with 4 % PFA for 10 min at RT. After several rinses with 0.1 % Triton X-100 in PBS (PBST), the tissue was incubated in HCl 2N solution (Fisher) for 30 min at 37 °C, then rinsed in borate buffer (0.1 M boric acid in distilled H_2_O, pH 8.5; Sigma-Aldrich) for 10 min and then rinsed several times with PBST. The sections were incubated for 1 h at RT in incubation buffer: 5% NDS diluted in PBST. Sections were then incubated overnight at 4 °C with sheep anti-BrdU (1:1,000; ab1893, Abcam) diluted in incubation buffer. The sections were kept for 30 min at RT and, after rinsing with PBST, antibody binding was detected with the corresponding fluorescent secondary antibody in incubation buffer for 1 h at RT (1:1,000, Invitrogen). The sections were rinsed again with PBST and PBS, and fixed with 4 % PFA for 10 min at RT. Then, the detection of Arg-I or EdU was performed as follows: Arg-I IHC was carried out by using a goat anti-Arg-I antibody (1:50; sc-18351, Santa Cruz Biotechnology) and EdU was revealed using Click-iT Alexa Fluor 488 Imaging Kit according to the manufactureŕs instructions (C110337, Invitrogen). The cell nuclei were stained with Hoechst 33342 (10 µg/mL, Sigma-Aldrich) and the sections were mounted with Fluoromount-G (Southern Biotech).

### Image acquisition and histopathological analysis of murine tissue

In all cases, 3 sections from each thoracic spinal cord (with a separation of 420 µm) were selected from each cohort of 12 EAE mice. A mosaic composition was obtained with 20X images of slices acquired on a confocal SP5 microscope (Leica) connected to a resonant scanning system at the Microscopy and Image Analysis Core Facility of the *Hospital Nacional de Parapléjicos*. Regions of interest corresponding to the demyelinated area or plaque, the adjacent periplaque and the NAWM were established mainly through the density of the cell nuclei within the infiltrated area. As such, the plaque of demyelinated lesions was characterized by a high nuclear density, while the periplaque was determined as the area corresponding to a 100 µm perimeter measured from the lesion edge (plaque) to the adjacent area. NAWM was established as the area corresponding to a perimeter of 100 µm measured from the edge to the periplaque to the deep white matter. Quantification of cell density was manually performed using Image J software.

### Statistical analysis

Data were expressed as mean ± SEM and analyzed with Sigma Plot version 11.0 (Systat Software Inc.). Shapiro-Wilk normality tests were performed on human and murine tissue samples. A Student’s t-test was used to compare different groups of mice (parametric statistical variables with a normal distribution) or Mann-Whitney U test was used for comparing non-parametric or non-normally distributed parametric variables. One-way analysis of variance (ANOVA) test or its corresponding ANOVA on ranks, followed by the properly *post-hoc* test, were carried out to compare between different areas of the lesions in mice or MS tissue. A Spearman test was carried out for the correlation analyses between disease duration and the ratio of OPC density (rAIL/cAIL) in MS tissue, as well as between peripheral blood M-MDSC content and the clinical signs in the EAE model. A Pearson test was used to correlate the density of OPC and the density of M-MDSCs in human samples, and the content of circulating M-MDSC and OPC/OL densities in the lesions of EAE mice. The minimum statistical significance was set at p < 0.05: ^$^p <0.05, ^$$^p <0.01, ^$$$^p <0.001, when comparing different animal groups; *p <0.05 when comparing rim and plaque or NAWM and plaque in human tissue, and periplaque or NAWM to the plaque in mice; ^#^p <0.05 when comparing plaque or NAWM to the periplaque in mice.

## List of abbreviations

Arg-I: Arginase-I
cAIL: core of mixed active/inactive lesions
CFA: complete Freund’s Adjuvant
CNS: central nervous system
DAB: 3, 3’-diaminobenzidine
EAE: experimental autoimmune encephalomyelitis
EC: eriochrome cyanine
EDSS: Expanded Disability Status Scale
FBS: fetal bovine serum
HLA: human leukocyte antigen
HPF: high power field
IHC: immunohistochemistry
IQR: interquartile range
ip: intraperitoneal
mAILs: mixed active/inactive lesions
MDSCs: myeloid-derived suppressor cells
M-MDSCs: monocytic-myeloid-derived suppressor cells
MOG: myelin oligodendrocyte glycoprotein
MRI: magnetic resonance imaging
MS: multiple sclerosis
NAWM: normal appearing white matter
NDS: normal donkey serum
OLs: mature oligodendrocytes
OPC: oligodendrocyte precursor cell
P/S: Penicillin/Streptomycin
PB: 0.1 M Phosphate Buffer, pH 7.4
PBS: sterile 1x Phosphate Buffered Saline
PBST: 0.1 % Triton X-100 in PBS
PET: positron emission tomography
PFA: paraformaldehyde
rAILs: rim of mixed active/inactive lesions
RRMS: relapsing-remitting multiple sclerosis
RT: room temperature
SI: severity index.

## DECLARATIONS

### Ethics approval and consent to participate

All the animal manipulations were approved by the institutional ethical committee (*Comité Ético de Experimentación Animal del Hospital Nacional de Parapléjicos*), and all the experiments were performed in compliance with the European guidelines for animal research (EC Council Directive 2010/63/EU, 90/219/EEC, Regulation No. 1946/2003), and with the Spanish National and Regional Guidelines for Animal Experimentation (RD 53/2013 and 178/2004, Ley 32/2007 and 9/2003, Decreto 320/2010). The study with human samples was approved by the local ethics committees of the *Complejo Hospitalario Universitario* de Toledo.

### Consent for publication

Not applicable.

### Availability of data and material

The datasets generated and/or analysed during the current study are available in the Zenodo repository (https://zenodo.org/records/12748820).

### Authors’ contributions

MPS-R: Investigation, Formal analysis, Writing - Original Draft, Visualization; CC-T: Investigation, Formal analysis, Visualization, Writing - Original Draft; IA-G: Investigation, Formal analysis; MCO: Investigation, Formal analysis, Writing - Original Draft, Visualization; IM-D: Investigation, RL-G: Investigation; JGA: Investigation; LC: Investigation; DC: Conceptualization, Resources, Writing - Original Draft, Writing - Review & Editing, Visualization, Supervision, Project administration, Funding acquisition. All authors have given approval to the final version of the manuscript.

## Acknowledgements

The authors would like to thank Dr Virginia Vila-del Sol and Ángela Marquina Rodríguez at the Flow Cytometry Core Facility of the *Hospital Nacional de Parapléjicos* and Dr José Ángel Rodríguez-Alfaro and Dr Javier Mazarío at the Microscopy Core Facility of the *Hospital Nacional de Parapléjicos* for their assistance with the flow cytometry analysis and the confocal imaging and histological quantifications.

